# Genetic Susceptibility to Enteric Fever in Experimentally Challenged Human Volunteers

**DOI:** 10.1101/2021.03.19.436030

**Authors:** Amber Barton, Jennifer Hill, Sagida Bibi, Liye Chen, Claire Jones, Elizabeth Jones, Susana Camara, Sonu Shrestha, Celina Jin, Malick M Gibani, Hazel Dobinson, Claire Waddington, Thomas C Darton, Christoph J Blohmke, Andrew J Pollard

**Author notes:** Corresponding author information, Amber J Barton.

## Abstract

**Background:** Infection with *Salmonella enterica* serovars Typhi and Paratyphi A cause an estimated 14 million cases of enteric fever annually. Here the controlled nature of challenge studies is exploited to identify genetic variants associated with enteric fever susceptibility.

**Methods:** Human challenge participants were genotyped by Illumina OmniExpress-24 BeadChip array (n=176) and/or transcriptionally profiled by RNA-sequencing (n=178).

**Results:** Two SNPs within *CAPN14* and *MIATNB* were identified with p<10^−5^ for association with development of symptoms or bacteraemia following oral *S*. Typhi or *S*. Paratyphi A challenge. Imputation of classical human leukocyte antigen (HLA) types from genomic and transcriptomic data identified HLA-B*27:05, previously associated with non-typhoidal *Salmonella*-induced reactive arthritis, as the HLA type most strongly associated with enteric fever susceptibility (p=0.012). Genes related to the unfolded protein response and heat shock were over-represented in HLA-B*27:05^+^ participants following challenge (p=0.01). Furthermore, intracellular replication of *S*. Typhi is higher in C1R cells transfected with HLA-B*27:05 (p=0.02).

**Conclusion:** These data suggest that activation of the unfolded protein response by HLA-B*27:05 misfolding may create an intracellular environment conducive to *S*. Typhi replication, increasing susceptibility to enteric fever.

## Background

*Salmonella enterica* serovars Typhi and Paratyphi A cause an estimated 14 million cases of enteric fever per year, resulting in 135,000 deaths [1]. Several risk factors have been identified for enteric fever, including poor sanitation and flooding [2]. Individual host factors also likely to contribute to disease susceptibility. Human challenge models, where volunteers are deliberately exposed to a pathogen, have been developed to study the biology of enteric fever and test experimental vaccines. Despite ingesting the same inoculation dose of bacteria, some challenged individuals remain infection-free, while others develop bacteraemia or symptoms consistent with enteric fever [3,4]. This could be explained in part by unmeasured factors such as effective bacterial dose reaching the intestinal mucosa, or other random effects not amenable to control. Alternatively, certain participants may have an innate resistance or susceptibility to enteric fever: in unvaccinated human challenge participants undergoing homologous re-challenge with S. Typhi, those who did not develop enteric fever on the first exposure were less likely to develop enteric fever on the second exposure [5]. Host genetics could play a role in this resistance. Genome-wide association studies (GWAS) are frequently used to find associations between genetic variants and complex non-Mendelian traits, with the aim of identifying genes which may provide insight into the pathology of a disease. For example, a GWAS identified polymorphisms in the NOD2 pathway as being associated with leprosy susceptibility [6]. NOD2 activation was later found to induce dendritic cell differentiation, which may protect against disease progression [7]. In the case of *Salmonella* infections, GWAS have revealed the HLA-DRB1*04:05 allele as conferring resistance against typhoid fever [8] and a locus in *STAT4* as being associated with non-typhoidal *Salmonella* bacteraemia [9].

In epidemiological studies, genetic heterogeneity in the pathogen is a confounder to the infected human host’s individual susceptibility to that pathogen, as illustrated by studies of tuberculosis, in which host SNPs predispose individuals to infection with a particular strain only [10]. In studies performed at our centre to date, only three strains of *Salmonella* have been used as a challenge agent, which has allowed us to statistically control for pathogen heterogeneity. All participants are exposed to the pathogen under highly controlled conditions, whereas in the field “non-infected controls” may have avoided infection due to lack of environmental exposure rather than having been exposed and resisted infection. Furthermore, prior exposure modifies susceptibility to enteric fever [5], which is difficult to account for in the field as *Salmonella* exposure is likely frequent during childhood in endemic settings. However this can be managed in challenge studies through strict inclusion criteria and careful screening, including exclusion of participants who had received a typhoid vaccine or lived in a typhoid-endemic area. Despite the advantages of human challenge studies, to our knowledge a GWAS has not previously been carried out on human challenge participants. Here we exploit this unique setting, and investigate how differences in host genetics relate to outcome of challenge. We identify SNPs within the genes *CAPN14* and *MIATNB* as having p<10^−5^ for association with development of enteric fever symptoms or bacteraemia following exposure. We find that HLA-B*27:05 is the HLA type most strongly associated with enteric fever susceptibility, enhancing intracellular replication of *S*. Typhi.

## Methods

### Enteric Fever Human Challenge Cohorts

Five enteric fever human challenge cohorts from studies conducted at the Centre for Clinical Vaccinology and Tropical Medicine (Churchill Hospital, Oxford, UK) were included in this analysis: a typhoid dose-finding study, a paratyphoid dose-finding study, a typhoid oral vaccine study, a typhoid Vi vaccine study, and a study investigating the role of the typhoid toxin, summarised in Table 1. All participants provided written informed consent. Following challenge, individuals with fever (sustained oral temperature ≥38°) or positive blood culture were diagnosed with enteric fever. All challenged participants were treated with ciprofloxacin or azithromycin either at time of diagnosis in diagnosed individuals, or after completing the 14-day challenge period if undiagnosed. Peripheral blood samples from participants from five different enteric fever human challenge cohorts were either genotyped or transcriptionally profiled, or in some cases both (Figure 1). A subset of participants underwent longitudinal transcriptional profiling, with data available from up to nine time points.

**Figure 1.**
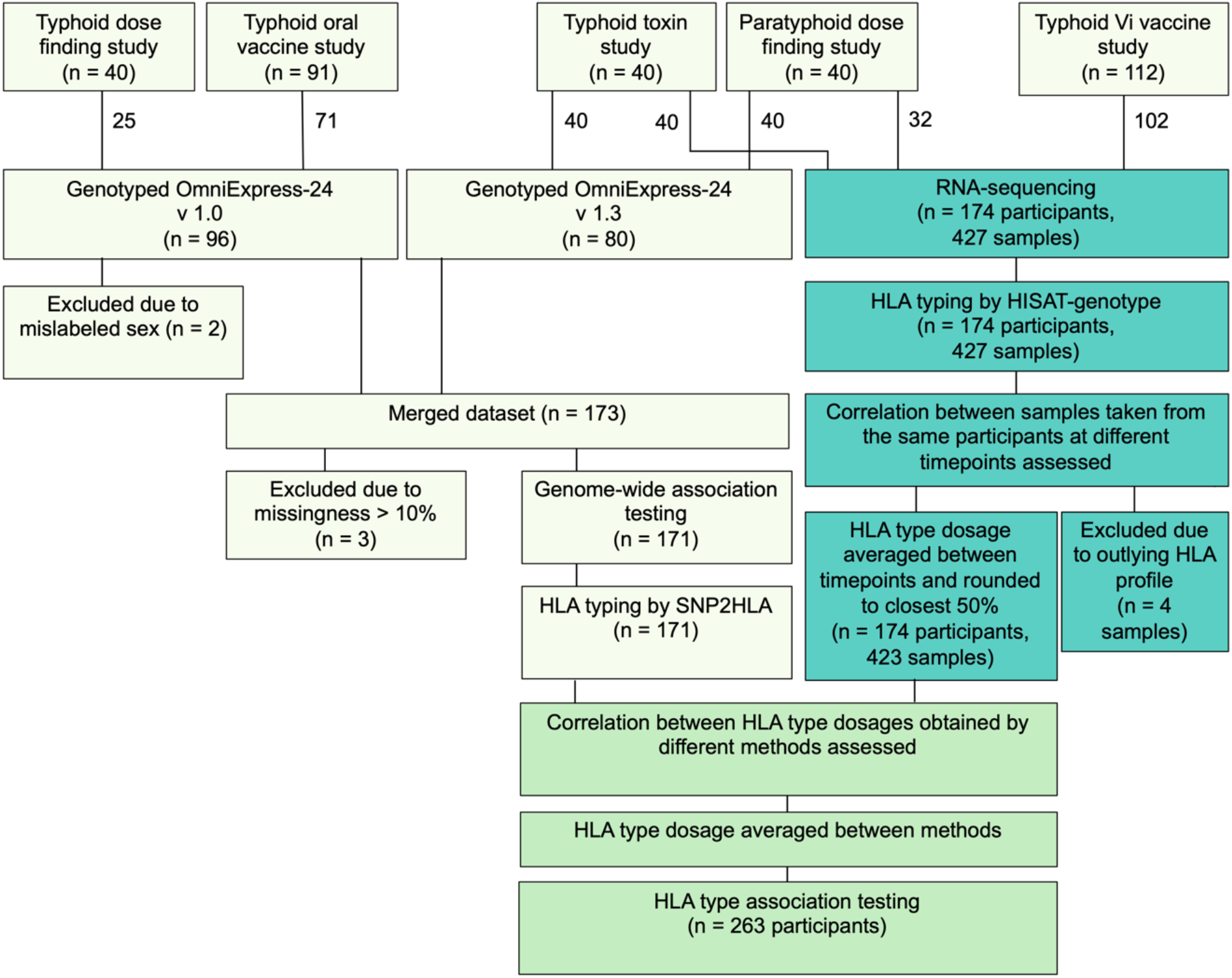
Number of participants and samples at each stage of the analysis pipeline

### Genotyping

DNA was extracted from blood clots using a QIA Symphony SP. Briefly, 180ul of ATL buffer (Qiagen™) was added to each clot and then vortexed and incubated overnight at 56°C for lysis. The following day 200ul of AL buffer (Qiagen™) was added to the lysed clot and mixed before transferring 500ul of the lysate to a 2ml tube and run on the QIA Symphony using the QIA Symphony DSP DNA Midi kit (Qiagen™). The protocol was a customised BC 400 protocol and DNA was eluted into 100ul. Samples were quantified using the Qubit and Qubit BR dsDNA reagents (Invitrogen). Samples from the typhoid dose finding and typhoid oral vaccine trial (total n=96) were genotyped by the Wellcome Trust Centre for Human Genetics using an Illumina OmniExpress-24 v1.0 BeadChip array, while samples from the paratyphoid dose finding study and typhoid toxin study (total n=80) were genotyped by Cambridge Genomic Services using an Illumina OmniExpress-24 v1.3 BeadChip array. Data cleaning for association analysis was carried out in PLINK [14]. Data processing steps are summarised in Figure 1.

Association analysis was carried out using a logistic regression model in PLINK, with challenge dose, vaccination status and principal components as covariates. The online tool SNPnexus [15] was used to identify genes proximal to SNPs. With the HapMap CEU dataset as a reference, SNP2HLA software [16] was used to impute single nucleotide polymorphisms in the HLA region and identify HLA alleles.

### RNA-sequencing

Whole blood samples were collected in Tempus Blood RNA tubes. RNA samples from the paratyphoid dose-finding and Vi vaccine trial were poly-A selected and underwent paired-end using a HiSeq V4 at the Wellcome Trust Sanger Institute. RNA samples from the typhoid toxin study underwent poly-A selection and paired end sequencing at the Beijing Genomics Institute using an Illumina HiSeq4000. Fastq files from the same sample were concatenated. Paired fastq files were aligned to a pre-built graph reference using HISAT2, followed by extraction of HLA-aligning reads. HLA typing and assembly was then carried out using HISAT-genotype [17].

### Correlation between HLA types imputed from different time points and by different methods

To assess the consistency of HISAT-genotype in imputing HLA-types, we compared estimated HLA type doses given for the same participant from whole blood samples collected at different timepoints. A Pearson correlation analysis was carried out on raw dosages for each pairwise comparison between timepoints to give a Pearson correlation coefficient (R) and p value for strength of association. We calculated both whether within a participant, dosage of each HLA type was consistent between timepoints, and within a HLA type, whether the dosage for each participant was consistent between timepoints.

To assess agreement between HLA types imputed by HISAT-genotype and SNP2HLA, for 71 participants where both genotyping and RNA-sequencing data were available, dosage was rounded for the closest 50%. As for certain participants HISAT-genotype results were available from RNA samples taken at different timepoints, the median was taken for each participant. A Pearson correlation analysis was carried out to compare estimated dosages given by HISAT-genotype and SNP2HLA. As above, we calculated both whether within a participant, the dosage of each HLA type was consistent between methods, and within a HLA type, whether the dosage for each participant was consistent between methods.

### Association between HLA type and outcome

Dosages were rounded to the nearest 50%, and for participants with multiple timepoints HLA-typed by HISAT-genotype, any timepoints with outlying dosages (Figure S1) were excluded and the median of the remaining timepoints taken. HLA types where there was no significant correlation (p>0.05) between timepoints were excluded. For those with both SNP2HLA and HISAT-genotype derived HLA-types the mean dosage was then taken. HLA type data from all cohorts were then combined. A logistic regression model was used to identify HLA types associated with outcome (1=diagnosed with enteric fever, 0=remained undiagnosed). Vaccination status, challenge dose and challenge strain were included as covariates. Statistical tests were carried out in R.

### Intracellular survival of *S*. Typhi in HLA-B*27:05^+^ cells

HLA-B*27:05^+^ C1R cells generated using lentiviral constructs were provided by the Bowness Group [18]. Transfected control and HLA-B*27:05^+^ cells were seeded in a 96 well plate at a density of 100,000 cells per well. A frozen glycerol stock of 5 × 10^8^ CFU/ml *S*. Typhi Quailes strain was thawed and washed twice with RPMI 1640 Media. Cells were inoculated at a multiplicity of infection (MOI) of 50 in triplicate. After one hour, gentamycin was added at a concentration of 200ug/ml. 24 hours post-inoculation cells were washed twice with RPMI then resuspended in 50ul 1% Triton-X100. After two minutes, lysates were serially diluted in PBS and plated onto tryptone soya agar. Colonies were counted following overnight incubation at 37^°^C. A one-tailed t-test was used to assess whether the number of colonies was higher in HLA-B*27:05^+^ cells.

### Differences in gene expression in those with HLA-B*27:05

Pre-alignment quality control on sequenced samples from the paratyphoid dose-finding study and Vi vaccine trial was carried out using FASTQC. As all files had high phred scores (>25) across their length, all were aligned to the human genome (GRCh38 Gencode version 26) using STAR-2.6.1c [19]. Total reads per sample ranged from 16-44 million. Reads per gene were counted using the STAR GeneCounts mode. Principal component analysis was used for outlier detection, with no samples excluded on this basis. Non-protein coding and haemoglobin subunit genes were excluded. Count tables were filtered to exclude genes with <1 count per million (cpm) in >31 samples (the number of baseline samples in control participants challenged with *S*. Typhi) and normalised using weighted trimmed mean of M-values scaling (edgeR). The count matrix was transformed using limma voom, and a linear regression model fitted with vaccination status, challenge strain, sequence pool and dose as covariates and participant ID as a blocking variable. At baseline and 12 hours post-challenge, differential gene expression analysis between those with and without a copy of HLA-B*27:05 was carried out, filtering to genes with average log_2_(expression)>0.

### Gene set enrichment analysis

Differences in gene expression between human challenge participants with and without a copy of HLA-B*27:05 were ranked by t-statistic at both baseline and 12 hours post-challenge. The entire ranked gene list, including non-significantly differentially expressed genes, were input into GSEA 4.1.0 software [20]. A custom gene set was created containing genes relating to the unfolded protein response and heat shock response (Table S1). An enrichment score reflecting the degree to which these genes were over-represented at the top of each ranked gene list was calculated. The p value of the enrichment score was then calculated by the GSEA 4.1.0 software using an empirical phenotype-based permutation test procedure [20].

## Results

### No SNPs were significantly associated with the outcome of challenge at the genome-wide level

A genome-wide association analysis was carried out on genotyped participants (103 cases of enteric fever, 68 controls following data cleaning) in order to identify any SNPs associated with development of fever, symptoms or bacteraemia following *S*. Typhi or *S*. Paratyphi A challenge (Figure 2). No SNPs reached genome-wide significance, with two SNPs within the genes *CAPN14* and *MIATNB* giving a p value below 1×10^−5^ (Figure S2).

**Figure 2.**
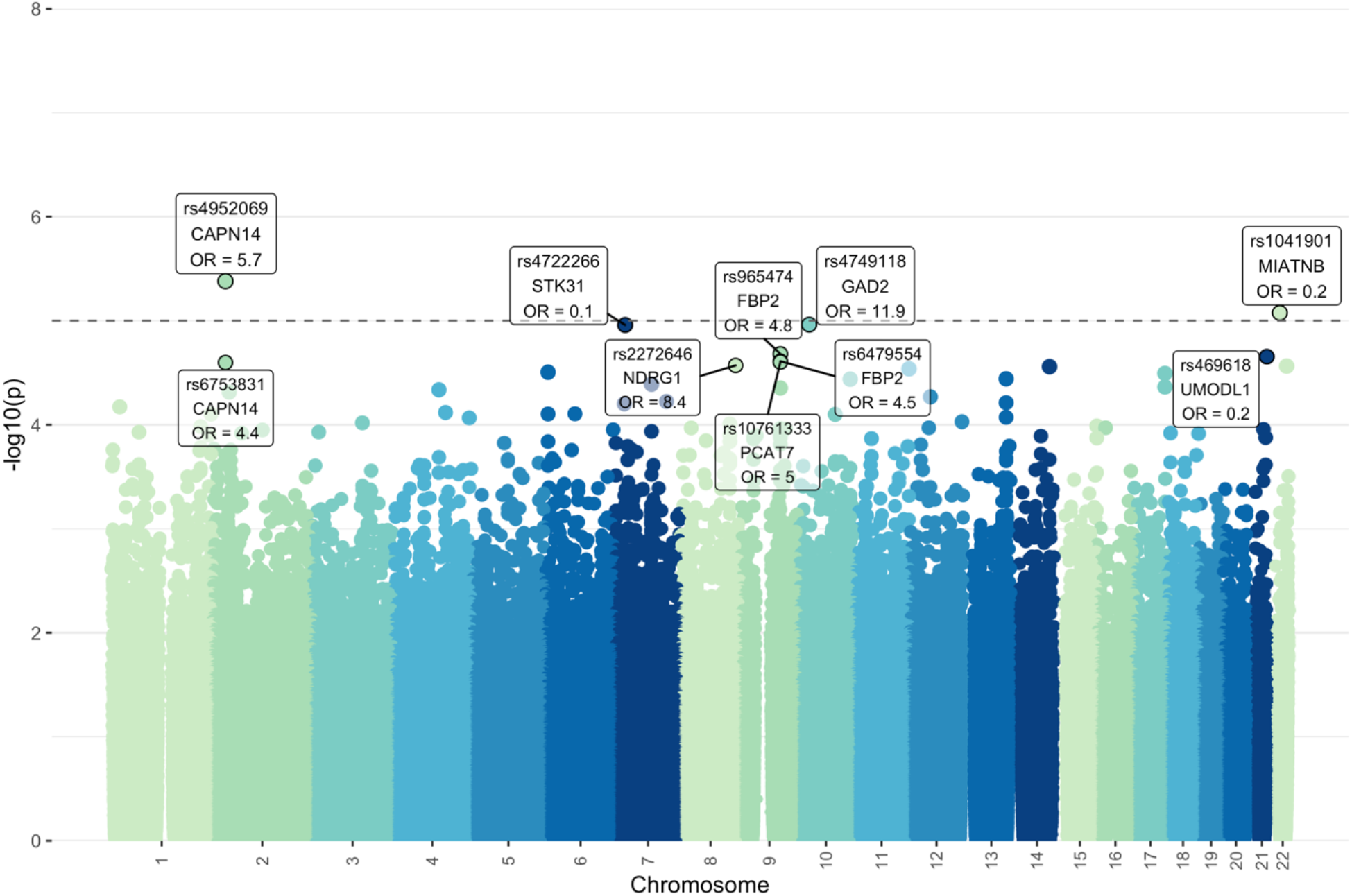
Manhattan plot showing the significance (-log_10_(unadjusted p value)) of the relationship between each single nucleotide polymorphism (SNP) and development of symptoms or bacteraemia following oral *S*. Typhi or *S*. Paratyphi A challenge, for each chromosome. The dotted line indicates a suggestive p value of 10^−5^. The ten SNPs with the lowest p values are highlighted, with the nearest proximal gene as identified by SNPnexus indicated as well as the odds ratio (OR).

### HLA-B*27:05 is associated with susceptibility to enteric fever

Given the number of individuals was too small to identify SNPs at the genome-wide level, we then focused on variation within the HLA region. HLA typing was performed either by imputation from genotyping data using SNP2HLA [16], or from raw RNA-sequencing data using HISAT-genotype [17]. We found HISAT-genotype to be highly consistent between RNA-sequencing samples taken from the same participant at different time points (Figures 3a and 3b; Table S3). For participants with both genotyping and RNA-sequencing data, HLA-typing using HISAT-genotype significantly correlated with the results of SNP2HLA imputation (Figures 3c and 3d).

**Figure 3:**
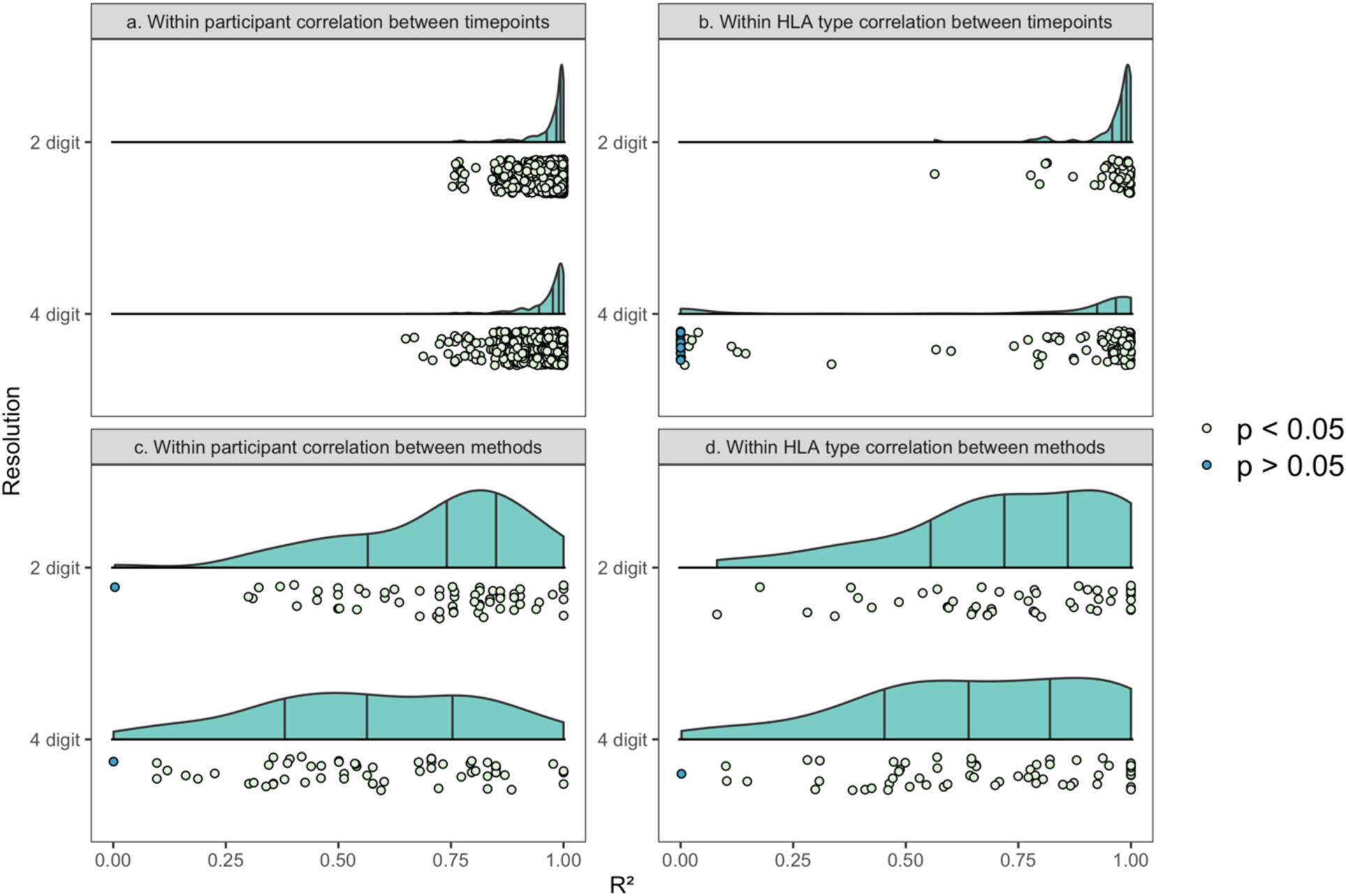
Distribution of squared Pearson correlation coefficients (R^2^) for HLA types at a 2-digit resolution and 4-digit resolution. Quantiles are indicated by vertical lines. Points are coloured by whether in a test of whether the Pearson correlation coefficient is different to zero, the p value was below 0.05. a. Correlation between the doses of each HLA type at different timepoints within each participant. HLA types were profiled from RNA-sequencing samples using HISAT-genotype, giving a dose (0-100%) of each HLA type for each participant. Each point represents one comparison; for participants where more than two timepoints were profiled, more than one point is shown per participant. b. Correlation between the doses for each participant at different timepoints within each HLA type. HLA types were profiled from RNA-sequencing samples using HISAT-genotype, giving a dose (0-100%) of each HLA type for each participant. Each point represents one HLA type. c. Correlation between the median doses of each HLA type for the same participant using either SNP2HLA imputation from genotyping data or HISAT-genotype typing from RNA-sequencing data. Each point represents one participant. d. Correlation within each HLA type between the doses for each participant using either SNP2HLA imputation from genotyping data or HISAT-genotype typing from RNA-sequencing data. Each point represents one HLA type.

The most common HLA-A, -B, -C, -DQA, -DQB1 and -DRB1 allele groups were A*02, B*07, C*07, DQA*01, DQB1*06 and DRB1*15 respectively (Figure 4a; Table S4). To identify whether any HLA types were associated with enteric fever, a logistic regression was carried out on HLA types at a 2-digit resolution. The HLA type most associated with susceptibility was HLA-B*27 (p=0.015, odds ratio=1.04, 95% confidence intervals 1.01-1.08, Figure 4b). While a small odds ratio, this finding was of particular interest as HLA-B*27 has been associated with non-typhoidal *Salmonella*-induced reactive arthritis and ankylosing spondylitis [21–24]. At 4-digit resolution this association was driven by HLA-B*27:05 (Figure 4c). Of 10 participants heterozygous for HLA-B*27:05, 9 were diagnosed with enteric fever (Figure 4d). While HLA-B*27:05 is most common in European populations, and the cohort analysed was predominantly white (Figure 5a), in the 1000 Genomes project HLA-B*27:05 was present in both the Punjabi population in Pakistan and Bengali population in Bangladesh, two countries where enteric fever is endemic [1,25](Figure 5b).

**Figure 4.**
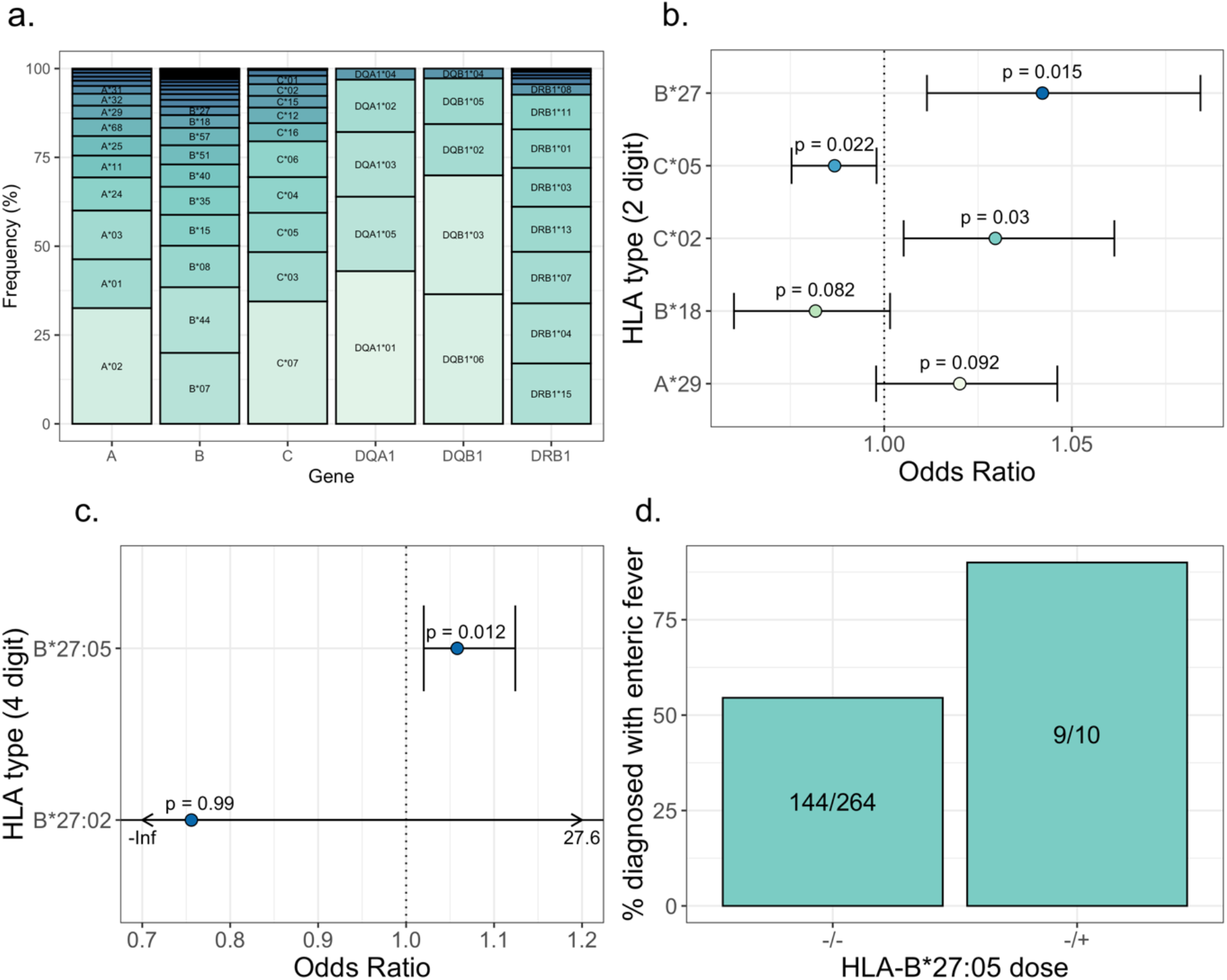
a. Relative frequency of each HLA type at a resolution of 2 digits for HLA-A, HLA-B, HLA-C, HLA-DQA1, HLA-DQB1 and HLA-DRB1 in the entire combined cohort, including participants from the typhoid dose finding study, typhoid oral vaccine trial, typhoid Vi vaccine trial, paratyphoid dose finding study and typhoid toxin study. b. Odds ratios (odds ratio >1 indicates associat]on with susceptibility and <1 with resistance) and 95% confidence intervals for the five HLA types most significantly associated with outcome of challenge at a resolution of 2 digits. P values are indicated for each. c. Odds ratios for the two HLA-B*27 sub-types at a resolution of 4-digits with 95% confidence intervals. P values are indicated for each. d. Percentage of participants who were diagnosed with enteric fever following challenge, stratified by the presence or absence of one copy of HLA-B*27:05. The proportion of participants diagnosed is indicated for each group.

**Figure 5.**
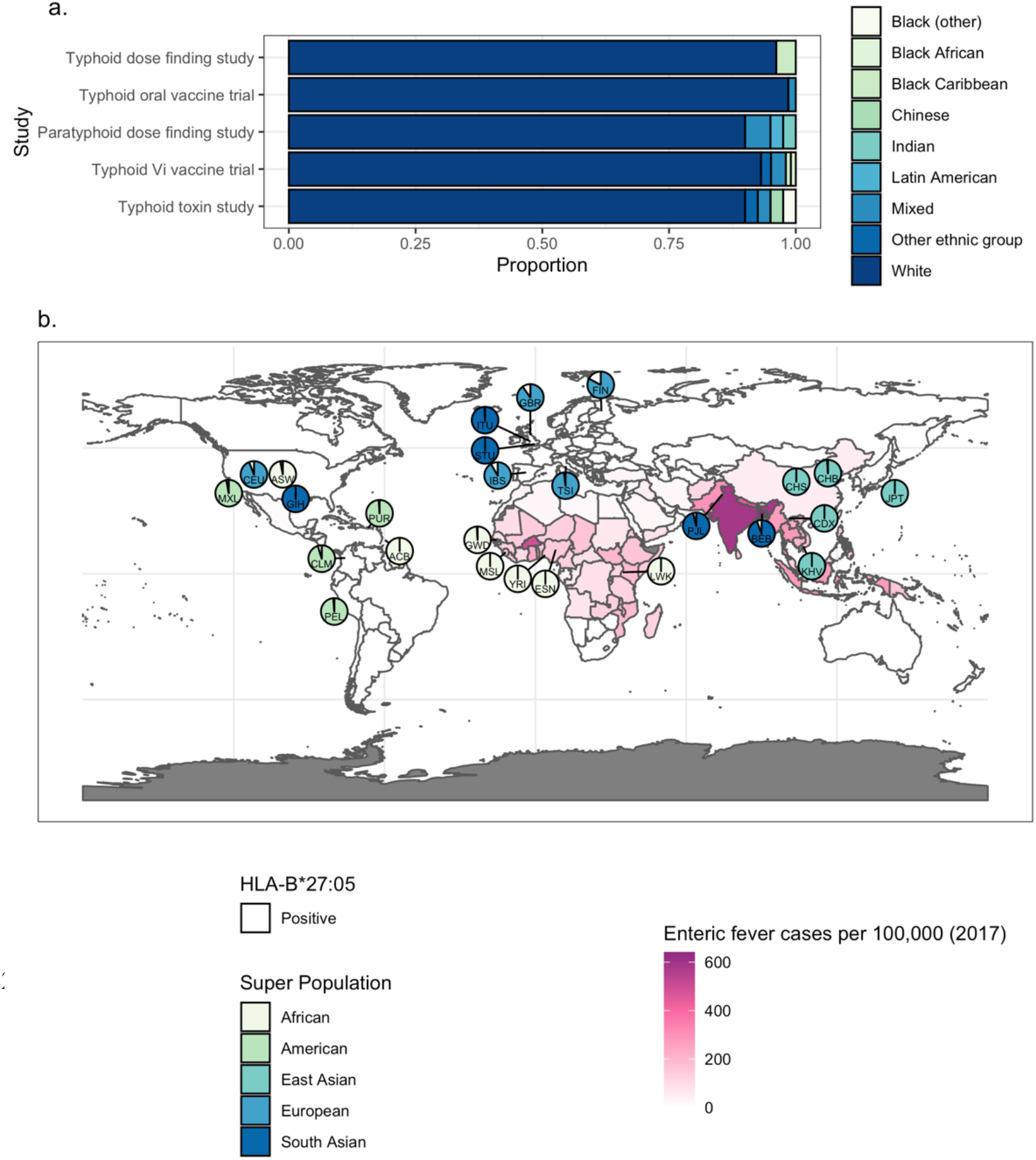
a. Self-reported ethnicity of participants within each study. b. The white sector of each pie chart indicates the proportion of 1000 Genome Project participants with at least one HLA-B*27:05 allele in each population. The remainder of the pie chart is coloured by ancestral continental region. Each country is coloured by enteric fever incidence rate per 100,000 as estimated by Stanaway et al. 2019. CHB = Han Chinese in Beijing, China, JPT = Japanese in Tokyo, Japan, CHS = Southern Han Chinese, CDX = Chinese Dai in Xishuangbanna, China, KHV = Kinh in Ho Chi Minh City, Vietnam, CEU = Utah Residents with Northern and Western European Ancestry, TSI = Toscani in Italia, FIN = Finnish in Finland, GBR = British in England and Scotland, IBS = Iberian Population in Spain, YRI = Yoruba in Ibadan, Nigeria, LWK = Luhya in Webuye, Kenya, GWD = Gambian in Western Divisions in the Gambia, MSL = Mende in Sierra Leone, ESN = Esan in Nigeria, ASW = Americans of African Ancestry in SW USA, ACB = African Caribbeans in Barbados, MXL = Mexican Ancestry from Los Angeles USA, PUR = Puerto Ricans from Puerto Rico, CLM = Colombians from Medellin, Colombia, PEL = Peruvians from Lima, Peru, GIH = Gujarati Indian from Houston, Texas, PJL = Punjabi from Lahore, Pakistan, BEB = Bengali from Bangladesh, STU = Sri Lankan Tamil from the UK, ITU = Indian Telugu from the UK.

To investigate the mechanism by which HLA-B*27:05 may contribute to enteric fever susceptibility, C1R cells transfected with HLA-B*27:05 were infected with *S*. Typhi in vitro for 24 hours. Compared with non-transfected controls, higher numbers of viable bacteria were recovered from HLA-B*27:05^+^ cells (Figure 6a), suggesting a mechanism independent of antigen presentation. This is consistent with previous literature finding that HLA-B*27:05 lowers the threshold for induction of the unfolded protein response, a pathway that is induced by and enhances intracellular *S*. Typhimurium infection [26].

**Figure 6.**
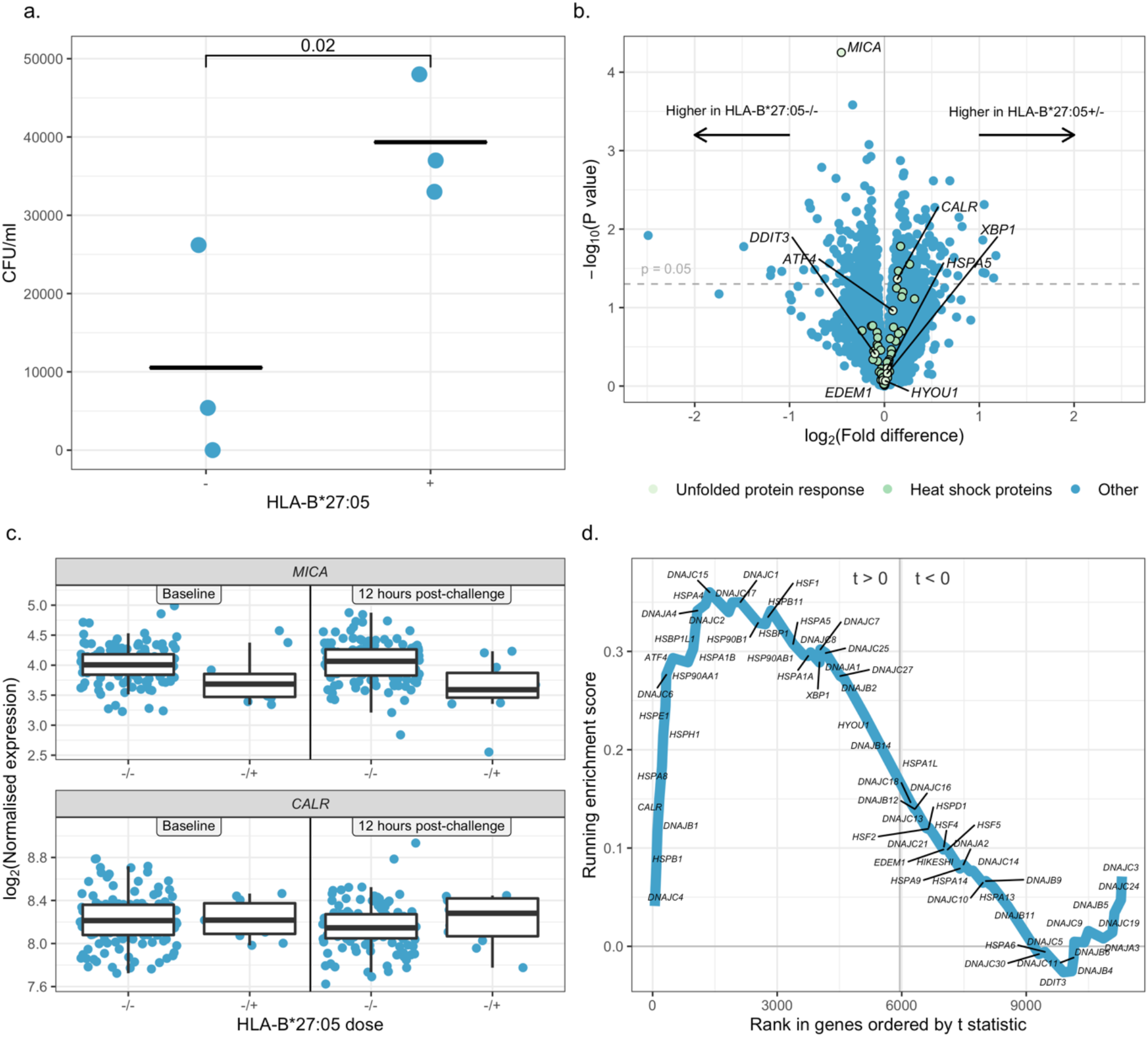
a. Colony forming units per ml recovered from C1R cells infected with S. Typhi Quailes strain, in the presence or absence of HLA-B*27 expression, 24 hours post-infection. Parent and HLA-B*27:05+ cells were seeded in a 96 well plate at a density of 100,000 cells per well, and infected with S. Typhi Quailes strain at an MOI of 0 or 10 in triplicate. After one hour gentamycin was added to kill extracellular bacteria. 24 hours post-inoculation cells were lysed using 1% Triton-X100, and lysates serially diluted and plated onto tryptone soya agar. Colonies were counted following overnight incubation at 37°C. A p value for a t-test is indicated. Points represent replicates within a single experiment. b. Volcano plot showing the log_2_(Fold Difference) in gene expression between HLA-B*27:05 positive and negative participants 12 hours post-challenge against the -log_10_(p-value). A dashed line indicating where p = 0.05 is shown, and genes relating to the unfolded protein response and heat shock proteins are highlighted. Genes more highly expressed in participants who were HLA-B*27:05 positive are shown positive further to the right, and those more highly expressed in HLA-B*27:05 negative participants further to the left. RNA expression was characterised by RNA-sequencing. Data were filtered, normalised and transformed, and differential expression then assessed using the limma R package, using participant ID, sequencing pool, vaccination status, challenge strain and dose as blocking variables. c. Expression of *MICA* and *CALR* following normalisation and transformation using the edgeR and limma packages, in HLA-B*27:05 positive and negative participants at baseline and 12 hours post-challenge. d. Running enrichment score for a custom gene set containing genes involved in the unfolded protein and heat shock response. Gene set enrichment analysis calculates an enrichment score by walking down a list of genes ranked by t statistic. When a gene within a gene set is encountered the running enrichment score increases, and when a gene outside the gene set is encountered it decreases. The enrichment score is the maximum deviation from zero. The genes in the custom gene set are indicated.

To investigate whether differences in the unfolded protein response can be detected in human challenge participants, we explored transcriptional differences between those who did and did not possess a copy of HLA-B*27:05 in the paratyphoid dose finding and Vi vaccine trial studies. We hypothesised that outcome of challenge is dependent on events occuring early after exposure and preceding development of acute disease, and therefore focused on 12 hours post-challenge, the timepoint at which dissemination of typhoidal *Salmonella* is thought to take place in the blood [27,28]. At 12 hours post-challenge with *S*. Typhi or *S*. Paratyphi A, the most significant differentially expressed gene expressed gene between the two groups was *MICA* (MHC Class I Polypeptide-Related Sequence A), encoding a ligand for NK cell activating receptor NKG2D (Figures 6b and 6c). Expression of *MICA* is inhibited by the unfolded protein response [29] and was expressed at lower levels by those with a copy of HLA-B*27:05 (p=0.00006 12 hours post-challenge, p=0.006 at baseline, linear modelling). The gene *CALR* encoding the calcium-binding chaperone calreticulin was more highly expressed in those with HLA-B*27:05 at 12 hours post-challenge but not at baseline (p=0.04 12 hours post-challenge, p=0.8 at baseline, Figure 6c). Gene set enrichment analysis [20] was then used to assess whether transcripts encoding proteins involved in the unfolded protein response were enriched amongst those with HLA-B*27:05. A custom gene set containing *CALR, ATF4, DDIT3, HSPA5, XBP1, EDEM1, HYOU1* as well as 61 genes annotated as relating to the heat-shock response was over-represented in those with HLA-B*27:05 at 12 hours post-challenge when ranked by t-statistic (p=0.01, Figure 6d). This gene set was not over-represented at baseline (p=0.4).

## Discussion

This study investigated genetic susceptibility to enteric fever in a human challenge setting. We found HISAT-genotype to be a consistent tool to impute HLA types from RNA-sequencing data, with HLA dosages correlating significantly with SNP2HLA dosages imputed from genotyping data. Of the HLA-types, HLA-B*27:05 was most associated with susceptibility to infection (p=0.012). Although participants were predominantly European, HLA-B*27:05 is also present in certain South Asian populations. HLA-B*27:05 mis-folds in the endoplasmic reticulum (ER), reducing the threshold for activation of the unfolded protein response, and has been linked with both reactive arthritis following Gram-negative bacterial infection [26], and ankylosing spondylitis [30]. When infected in vitro with *S*. Typhimurium, both monocyte-like U937 cells and epithelial HeLa cells transfected with HLA-B*27:05 exhibit higher levels of intracellular replication [26,31]. Although the exact mechanism is unknown, the unfolded protein response appears to create a favourable environment for *S*. Typhimurium, the presence of HLA-B*27:05 increasing its expression of SPI-2 genes [32] and causing it to replicate at the cell periphery [26]. Enhanced replication of *S*. Typhimurium was abrogated when HLA-B*27:05 was stabilised by fusion with beta-2-microglobulin [26]. Pharmacological induction of ER stress by thapsigargin enhances *S*. Typhimurium replication, while infection with *S*. Typhimurium stimulates the unfolded protein response by a mechanism dependent on bacterial effector *sifA* [26]. Although *sifA* is also present in *S*. Typhi, its sequence differs to sifA in *S*. Typhimurium [33]. However, we still observed enhanced replication of *S*. Typhi Quailes strain in C1R cells (p=0.02, one-tailed t-test), suggesting this phenomenon is not serovar-specific.

The gene encoding ER chaperone calreticulin, *CALR*, was higher in HLA-B*27:05^+^ human volunteers 12 hours following enteric fever challenge but not at baseline (p=0.04 12 hours post-challenge, p=0.8 at baseline). Gene set enrichment analysis [20] was then used to assess whether transcripts encoding proteins involved in the unfolded protein response were enriched amongst those with HLA-B*27:05. A custom gene set containing genes annotated as relating to the unfolded protein response and heat-shock response was over-represented in those with HLA-B*27:05 at 12 hours post-challenge when ranked by t-statistic (p=0.01). However this gene set was not over-represented at baseline (p=0.4). This supports the hypothesis that HLA-B*27:05 reduces the threshold for unfolded protein response activation in infection. At 12 hours post-challenge, the most significant differentially expressed gene expressed gene between the two groups was *MICA*, encoding a ligand for NK cell activating receptor NKG2D (p=0.00006, linear modelling). *MICA* is downregulated by the unfolded protein response [29], and was expressed at lower levels in participants with HLA-B*27:05 12 hours post-challenge. In viral infections, downregulation of *MICA* prevents recognition by NK cells [34]. Polymorphisms in *MICA* have been related to susceptibility to leprosy, which, in common with enteric fever, infects mononuclear phagocytes [35–37]. In contrast to *CALR, MICA* was also differentially expressed in HLA-B*27:05^+^ participants at baseline (p=0.06, linear modelling), suggesting either that HLA-B*27:05 can induce certain aspects of the unfolded protein response in the absence of infection, or that its decreased expression is mediated by a different mechanism.

In the absence of SNPs with very high odds ratios in our cohort, we were underpowered to detect significant SNPs at a genome wide level. The SNP with the lowest p value (rs4952069, 4.2 × 10^−6^) falls in the intronic region of *CAPN14*, a calcium-dependent cysteine protease regulated by Th2 cytokines IL-13 and IL-4 [38]. Intronic SNPs may either be linked to a causative coding SNP, or themselves affect gene expression through splicing or transcription factor binding [39]. *CAPN14* is thought to play a regulatory role in the oesophageal epithelium, with overexpression impairing barrier function, and SNPs in this locus having been associated with susceptibility to the allergic inflammatory disease eosinophilic oesophagitis [40] and middle ear infection [41]. While the cellular response to enteric fever infection is Th1 dominated, Th2 cytokines may be modulated by infection, with *S*. Typhi-specific IL-13 secretion observed in peripheral blood mononuclear cells isolated during typhoid fever convalescence [45] and IL-4 secreted at the apical side of intestinal biopsies infected in vitro with *S*. Typhi [46]. Co-infection of mice with both *S*. Typhimurium and Th2-inducing hookworms impairs clearance of *S*. Typhimurium, suggesting that polarisation towards a Th2 response could be detrimental [47]. Therefore genetic variations predisposing individuals to a more Th2-dominant response to infection could feasibly affect susceptibility to enteric fever.

This is the first genetic study to investigate susceptibility to infection using samples obtained from human challenge volunteers. Furthermore, while HLA-B*27:05 has been linked to non-typhoidal *Salmonella* infections, this is the first study to find an association with enteric fever. However, we were limited by several factors. Firstly, there were cases where the HLA type of a participant was ambiguous, predominantly due to SNP2HLA suggesting several possible HLA types, but also incomplete agreement between SNP2HLA and HISAT-genotype dosages. Secondly, due to the nature of human challenge studies, our sample size was smaller than conventional GWAS. While notable GWAS with smaller samples than ours have included those associating genetic variants with vitiligo and response to anti-TNF treatment, a larger sample would have enabled us to detect associations with smaller effect sizes [48,49]. Only 10 participants were unambiguously identified as HLA-B*27:05^+^ and the odds ratio was small in magnitude, suggesting that HLA-B*27:05 explains only a small proportion of innate susceptibility to enteric fever. However, given previous evidence of an association with both *Salmonella-*induced reactive arthritis and intracellular *S*. Typhimurium replication, this intriguing association warrants validation by further studies. Finally, this study was carried out a predominantly European cohort not previously exposed to typhoidal *Salmonella*. While this allowed us to investigate genetic susceptibility without the confounding factor of previous exposure, it is not representative of the population in an endemic setting. However, it could have implications for travel medicine: for example, those with a family history of ankylosing spondylitis could be strongly encouraged to undergo typhoid vaccination prior to travelling. Although reactive arthritis following live oral typhoid vaccination is a rare complication [50], parenteral vaccination may be preferable in this case. Furthermore, HLA-B*27:05 is present both in Punjabi and Bengali populations, suggesting this allele could play a role in an endemic setting [25].

## Acknowledgements

The authors would like to thank the volunteers for participating in the studies, and the High-Throughput Genomics Group at the Wellcome Trust Centre for Human Genetics for the generation of genotyping data. We would also like to thank the Data Safety and Monitoring Committee for providing safety oversight of the studies, the University of Maryland for providing *S*. Typhi Quailes challenge strain and GSK Vaccines Institute for Global Health for providing *S*. Paratyphi A NVGH308 strain. Finally, we would like to acknowledge Paul Bowness for supervising transfection of the transgenic C1R cells, and the additional clinical and laboratory support provided by Oxford Vaccine Group staff during the enteric fever studies.

## Footnotes

### Meetings where work was previously presented

BSI virtual conference, December 2020, United Kingdom, Abstract ID 989

